# Gene flow and species delimitation in fishes of Western North America: Flannelmouth (*Catostomus latipinnis*) and Bluehead sucker (*C. Pantosteus discobolus*)

**DOI:** 10.1101/763532

**Authors:** Max R. Bangs, Marlis R. Douglas, Tyler K. Chafin, Michael E. Douglas

## Abstract

The delimitation of species-boundaries, particularly those obscured by reticulation, is a critical step in contemporary biodiversity assessment. It is especially relevant for conservation and management of indigenous fishes in western North America, represented herein by two species with dissimilar life-histories co-distributed in the highly modified Colorado River (i.e., Flannelmouth Sucker, *Catostomus latipinnis*; Bluehead Sucker, *C. Pantosteus discobolus*). To quantify phylogenomic patterns and examine proposed taxonomic revisions, we first employed double-digest restriction-site associated DNA sequencing (ddRAD), yielding 39,755 unlinked SNPs across 139 samples. These were subsequently evaluated with multiple analytical approaches and by contrasting life history data. Three phylogenetic methods and a Bayesian assignment test highlighted similar phylogenomic patterns in each, but with considerable difference in presumed times of divergence. Three lineages were detected in Bluehead Sucker, supporting elevation of *C. P. virescens* to species-status, and recognizing *C. P. discobolus yarrowi* (Zuni Bluehead Sucker) as a discrete entity. Admixture in the latter necessitated a reevaluation of its contemporary and historic distributions, underscoring how biodiversity identification can be confounded by complex evolutionary histories. In addition, we defined three separate Flannelmouth Sucker lineages as ESUs (Evolutionarily Significant Units), given limited phenotypic and genetic differentiation, contemporary isolation, and lack of concordance (per the genealogical concordance component of the phylogenetic species concept). Introgression was diagnosed in both species, with the Little Colorado and Virgin rivers in particular. Our diagnostic methods, and the alignment of our SNPs with previous morphological, enzymatic, and mitochondrial work, allowed us to partition complex evolutionary histories into requisite components, such as isolation *versus* secondary contact.

## 1 INTRODUCTION

The delimitation of species (i.e., the process by which boundaries are not only identified but new species discovered; Wiens, 2007), is a fundamental issue in biology, and its mechanics contain aspects both theoretical and applied (Carstens et al., 2013). It is a requirement not only for effective biodiversity conservation (Frankham, 2010) but also for management, particularly with regards to the Endangered Species Act (ESA) (Waples, 1991). However, distinct boundaries traditionally assumed to characterize species (Sloan, 2008, Baum, 2009), are particularly difficult to identify early in the speciation process (Sullivan et al., 2013) or within groups where extensive reticulation has occurred (Mallet et al., 2015). This has led to an evolving interpretation of the speciation process, now viewed as a continuum from population through various ascending steps, but with genealogical distinctiveness achieved gradually and manifested differentially across the genome (Mallet, 2001).

Another complicating factor is the daunting number of narrow concepts that now encapsulate speciation (Coyne and Orr, 2004). Here, it is important to recognize that species are defined by morphological and genetic gaps, rather than a define ‘process’ and thus contribute but little to species delineation. Clearly, a more nuanced approach (see below) would help define conservation units across the population-species continuum. This would not only advance species conservation but also promote effective management strategies to protect genetic diversity (as stipulated in Strategic Goal C of the 2010 United Nations Convention on Biological Diversity < https://www.cbd.int/sp/targets/ >).

The most commonly utilized approach for delineating lineages is DNA-based, but with questionable reliance upon a single marker (i.e. DNA barcoding; Ahrens et al., 2016), especially given the semipermeable boundaries now recognized in species. Despite being cofounded by various problems, single-gene methods still predominate in the literature, often with sample sizes that do not capture intraspecific haplotype variability, an issue scarcely parameterized (Phillips et al., 2019). Additionally, single-locus delimitation methods fall under several broad categories, yet each suffers from limitations not easily overcome (Dellicour & Flot, 2018).

However, genomic DNA techniques are increasingly being applied to more formally delineate lineages (Allendorf et al., 2017). SNP (single nucleotide polymorphisms) panels are being applied to not only broaden and extend signals of population and species differentiation, but also to unravel their interrelationships. Two insights have emerged so far from the application of contemporary genomic approaches. First increased resolution provided by SNP panels has identified populations that diverge markedly within taxa previously-identified. These are subsequently characterized as ‘cryptic’ species (Singhal et al., 2018; Spriggs et al., 2019), with population histories not only statistically inferred but also tested against alternative models of divergence. A second insight is the revelation that admixture among lineages is not only quite common but also greater than previously thought (Dasmahapatra et al., 2012; Fontaine et al., 2015; Quattrini et al., 2019), even to the extent of promoting new adaptive radiations (Lamichhaney et al., 2018).

The increased resolution provided by reduced-representation genomic approaches has negative connotations as well. For example, elevated lineage resolution (as above) has the propensity to re-ignite earlier debates regarding the over-splitting of species (Isaac et al., 2004; Sullivan et al., 2014). Not surprisingly, this process is rife with value judgments, one of which seemingly intuits that species defined by parsing previously identified biodiversity are more problematic than those discovered *de novo* (Padial and de la Riva, 2006; Sullivan et al., 2014). Similar issues emerge when intraspecific diversity is interpreted for conservation actions (Funk et al., 2012). For example, the Evolutionarily Significant Unit (ESU) was conceived as a complement to existing taxonomy (Ryder, 1986; Moritz, 1994), with an intent to quickly identify conservation units worthy of protection without resorting to a laboriously slow and unwieldy taxonomic categorization. Again (as above), population genomics provides an abundance of neutral loci for delimitation of ESUs. However, despite its intended simplicity, the ESU concept (Frazer & Bernatchez, 2001; Holycross & Douglas, 2007) has seemingly become an either/ or categorization (i.e., dichotomized) such that it not only contravenes the continuum through which populations evolve, but also reflects those difficulties that emerge when subspecies are designated arbitrarily from continuous geographic distributions (Douglas et al., 2006).

Similarly, the Management Unit (MU) is yet another conservation category with traditional roots, generally referred to in fisheries literature as a ‘stock’ (Ryman & Utter, 1986). It now has a more contemporaneous meaning, defined primarily by population genomic data, and represents a conservation unit isolated demographically from other such units (Palsbøll et al., 2014; Mussmann et al., 2019). While genomic data clearly hold great potential for elucidating the evolutionary process, arguments must still be resolved before they become a de facto diagnostic tool for species delineation (Stanton et al., 2019). For example, genomic techniques were unsuccessful in unravelling a hybrid complex among Darwin’s Finches in the Galapagos (Zink & Vázguez-Miranda, 2019).

In this study, we applied a contemporary framework for genomic analysis (Leaché & Fujita, 2010), by initially clustering our admixed study species so as to detect erroneous species-designations derived from inter-specific gene flow (Camargo et al., 2012; Stewart et al., 2014). This approach gains additional power when multiple lines of evidence are integrated, such as life history, geographic distributions, and morphology (Knowles & Carstens, 2007; Schlick-Steiner et al., 2010; Fujita et al., 2012). As a result, the complex histories of study species can be more clearly discerned despite difficulties imposed by introgression. This is particularly appealing as herein, when problematic species are a focus of conservation concern (Pyron et al., 2016).

### 1.1 The Biogeography of our Study Species

Flannelmouth Sucker (*Catostomus latipinnis*) and Bluehead Sucker (*C. Pantosteus discobolus*) have complex evolutionary histories that reflect historical introgression (Smith et al., 2013), as well as contemporary hybridization with various congeners (Douglas & Douglas, 2010; Mandeville et al., 2015; Bangs et al., 2017). Both have been relatively understudied, yet their conservation concerns have accelerated due to a prolonged drought in western North America superimposed onto an ever-increasing anthropogenic demand for water (Seager & Vecchi, 2010). Given this, a federal and multi-state effort has now coalesced on basin-wide mitigation and recovery of both species (Carmen, 2007). Consequently, the accurate delimitation of species, as well as the designation of potential conservation units are highly relevant, especially given that our study species comprise, in an historic sense, the greatest endemic fish abundance/ biomass in the Upper Colorado River Basin (Hubbs et al., 1948).

Each species exhibits a different life history (Sigler and Miller, 1963; Behn & Baxter, 2019), with Flannelmouth Sucker primarily inhabiting the mainstem (Douglas et al., 1998, 2003) and Bluehead Sucker preferring higher elevation streams that have subsequently become more fragmented over time (Hopken et al., 2013). However, mark–recapture studies (Fraser et al., 2017) still emphasize tributaries in the upper Colorado River basin as important habitat for both species.

The response of our study species to the geologic history of western North America is an aspect of their life histories (reviewed in Bezzerides & Bestgen, 2002). Vicariant processes (i.e., vulcanism and drainage rearrangements) coupled with episodic drought, have induced long periods of isolation sporadically augmented by more pluvial periods that, in turn, have promoted secondary contact (Smith et al., 2010). Thus, a comparative study of each species can not only provide insights into the manner by which admixture has influenced their evolution, but also clarify our understanding of the Colorado River Basin itself.

Both species are primarily endemic to the Upper Colorado River Basin, but Flannelmouth Sucker is also found in the Virgin River of the Lower Colorado River Basin, and Bluehead Sucker in the neighboring Bonneville Basin to the west (Figure 1). The latter may potentially represent a different species (*C. P. virescens*), as judged by morphological (Smith et al., 2013), mitochondrial (Hopken et al., 2013; Unmack et al., 2014), and nuclear phylogenies (Bangs et al. 2018b). Taxonomic uncertainties within Flannelmouth and Bluehead sucker present additional management complications, especially with regard to the presence of potentially unique lineages in one major tributary - the Little Colorado River (Figure 2). One lineage may represent a unique species (i.e., Little Colorado River Sucker), currently grouped with Flannelmouth Sucker, whereas a second may be a unique subspecies (i.e., Zuni Bluehead Sucker, *C. P. d. yarrowi*) now found only in the Zuni River (NM) and Kin Lee Chee Creek (AZ), but with a presumed historic distribution that potentially included the entire Little Colorado River (Minckley, 1973).

**Figure 1:**
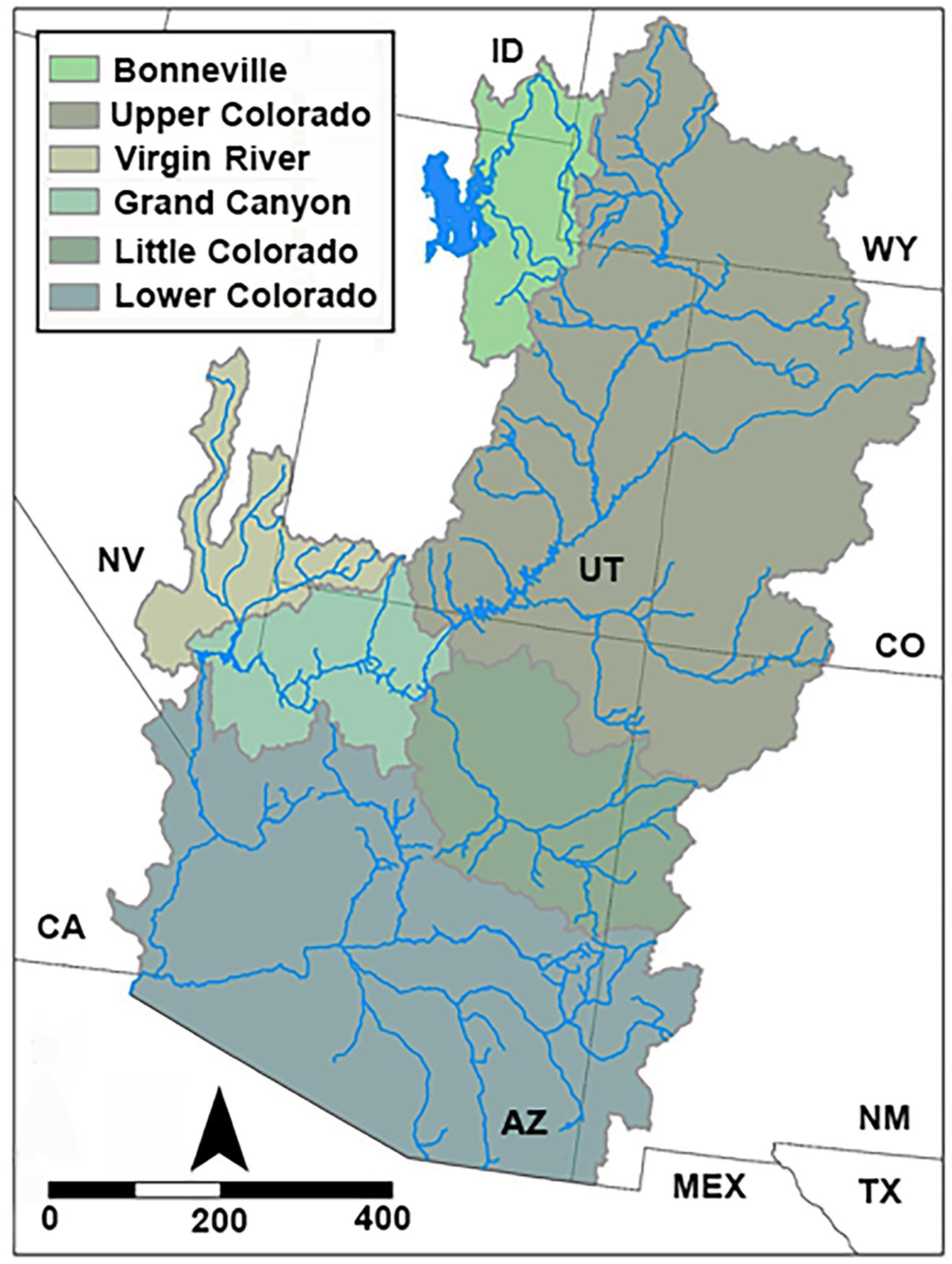
Map depicting the Colorado River and Bonneville basins, with adjacent basins or recognized geographic regions; ID=Idaho; MEX=México; NM=New Mexico; NV=Nevada; TX=Texas; UT=Utah; WY=Wyoming.

**Figure 2:**
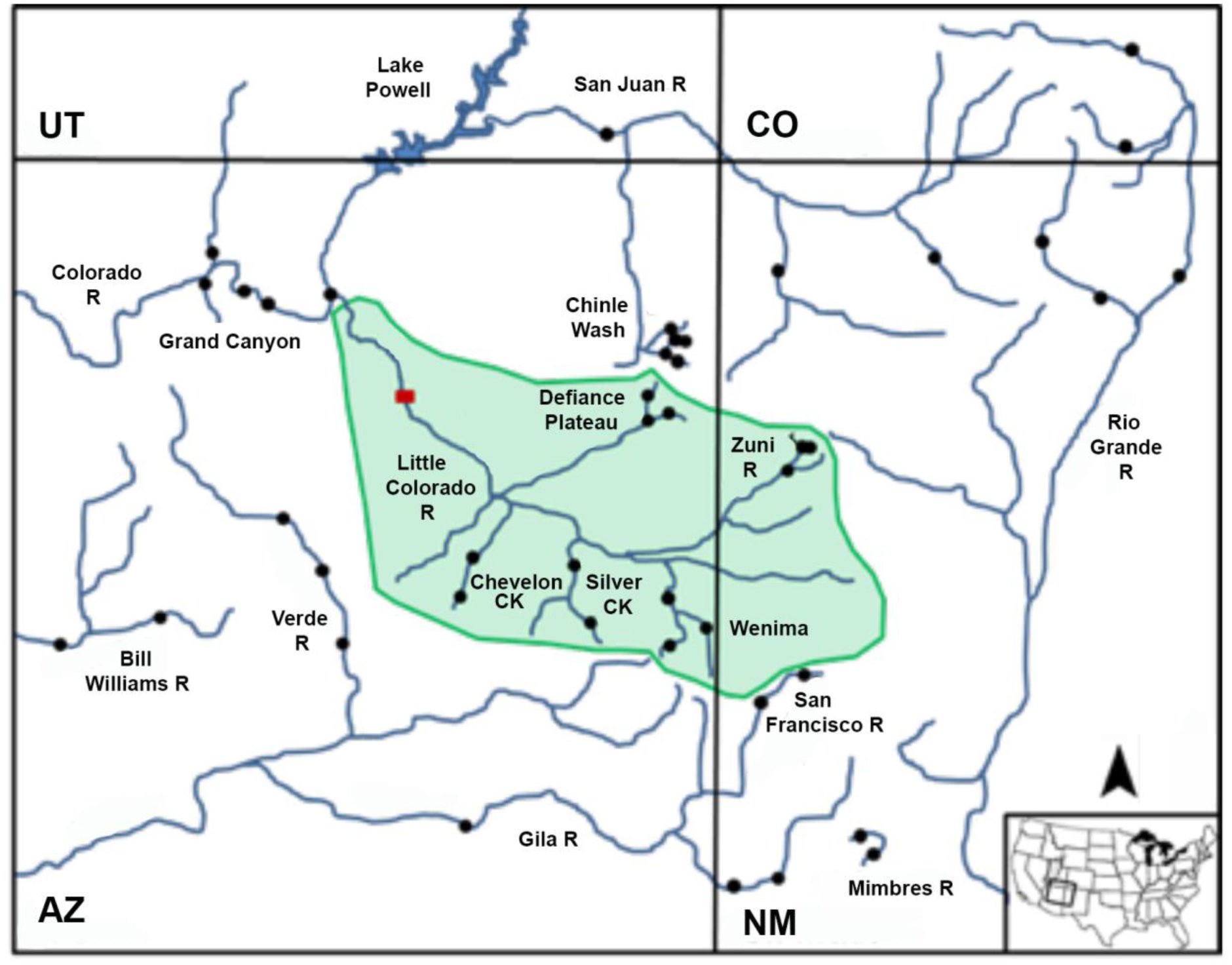
Topographic location of the Little Colorado River watershed (green) and surrounding drainages. Black dots represent collection sites. Red rectangle depicts Grand Falls, a vicariant barrier. AZ=Arizona; CO=Colorado; NM=New Mexico; UT=Utah.

The quantification of molecular variability in both catostomids is a key element in delimiting species-boundaries, management units, and historic patterns of reticulation. Here we build upon previous work (Bangs et al., 2018b) by applying species delimitation methods, phylogenomic (i.e., concatenated and multispecies coalescence), and population genomic approaches (i.e., Bayesian clustering and hybrid detection) to identify potential species range-wide, but with special focus on the Little Colorado River. In this regard, the impacts of divergent life histories, as well as the role of stream capture and hybridization, are particularly germane with regard to the breadth and depth of differentiation found within each.

## 2 METHODS

### 2.1 Sample acquisition

Samples were collected as either fin clips or tissue plugs between 1995 and 2011. Genomic DNA was extracted using the PureGene® Purification Kit or DNeasy® Tissue Kit (Qiagen Inc., Valencia CA) following manufacturer’s protocols, and stored in DNA hydrating solution. Additional samples were obtained from the Museum of Southwestern Biology (University of New Mexico) (see Acknowledgements).

A total of 139 samples (Table 1) included 81 (subgenus *Pantosteus*) and 57 (subgenus *Catostomus*) (per Smith et al., 2013). Bluehead Sucker samples (*C. P. discobolus*, N=65) spanned its range, including the Bonneville Basin (N=5), Grand Canyon AZ (N=10), Chinle Wash NM (N=10), Little Colorado River (N=29), and various sites in the Upper Colorado River Basin above Grand Canyon (N=11) (Figure 1, Table 1). Rio Grande Sucker (*C. P. plebeius*; N=6) and Desert Sucker (*C. P. clarkii*; N=8) were also examined so as to evaluate their potential contributions with regard to hybridization with other *Pantosteus*. Mountain Sucker from the Missouri River Basin (*C. P. jordani,* N=2) was included as outgroup for *Pantosteus*.

**Table 1:**
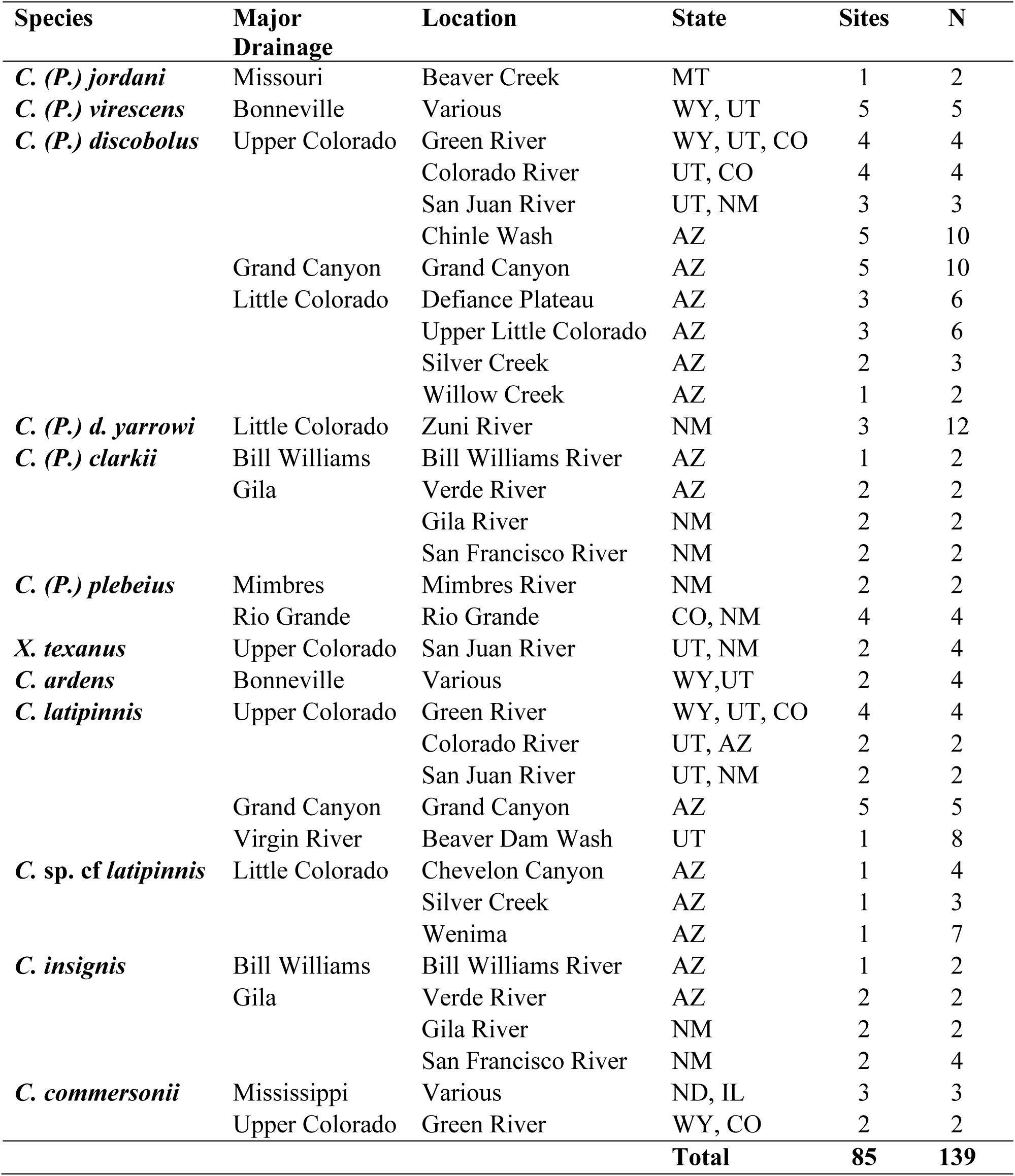
Sample sizes for each species by major drainage, location, and state. Included for each species are number of sample sites (Sites) and individuals (N).

Flannelmouth Sucker (N=35) was collected range-wide, to include the Virgin River UT (N=8), Little Colorado River (N=14), Grand Canyon AZ (N=5), and various sites in the Upper Colorado River Basin above Grand Canyon (N=8) (Figure 1). White Sucker (*C. commersonii*) obtained from locations in its native range (N=3), and an introduced population in the Colorado River (N=2), were incorporated as a *Catostomus* outgroup (Table 1). We also incorporated Sonora Sucker (*C. insignis*; N=10), Utah Sucker (*C. ardens*; N=4), and Razorback Sucker (*Xyrauchen texanus*; N=4) due to their geographic proximity, close phylogenetic relationships, and potential for hybridization with Flannelmouth.

### 2.2 Data collection

DNA was extracted with PureGene® Purification Kit or DNeasy® Tissue Kit (Qiagen Inc, Valencia CA) and stored in DNA hydrating solution (same kits). Libraries for double digest restriction-site associated DNA (ddRAD) were generated following Bangs et al. (2018b). This included: Digestion with PstI (5’-CTGCAG-3’) and MspI (5’-CCGG-3’), pooling 48 individuals prior to a size selection of 350-400bps, PCR amplification, and the combination of two libraries per lane of Illumina HiSeq, 2000 single-end 100bp sequencing. Samples for each reference species and region were randomly distributed across several libraries and lanes so as to reduce the potential for library preparation bias. Sequencing was performed at the University of Wisconsin Biotechnology Center (Madison).

### 2.3 Filtering and alignment

Illumina reads were filtered and aligned per Bangs et al. (2018b) using PYRAD v.3.0.5 (Eaton & Ree, 2013). This included: Clustering at a threshold of 80% based the uncorrected sequence variation in catostomid fishes (Chen & Mayden, 2012; Bangs et al., 2017), and removal of restriction site sequence and barcode. In addition, loci were removed if they displayed: 1) <5 reads per individual), 2) >10 heterozygous sites within a consensus, 3) >2 haplotypes for an individual, 4) >75% heterozygosity for a site among individuals, and 5) <50% of individuals at a given locus.

### 2.4 Clustering algorithm and phylogenetic methods

All analyses utilized the unlinked SNPs file generated from PYRAD, save for the concentrated SNP phylogenetic methods that required the all SNPs file. Bayesian clustering (STRUCTURE v. 2.3.4; Pritchard et al., 2000) employed the admixture model with correlated allele frequencies and a burn-in of 100,000 generation, followed by 500,000 iterations post-burn-in. No population priors were used. Genetic clusters (k=1-16) were each performed with 15 iterations, averaged across iterations to determine final values. The most likely clusters were resolved by using the estimated log probability of data Pr(x|k), and the Δk statistic (per Evanno et al., 2005). Bayesian clustering also substantiated that all contemporary hybrids with invasive White Sucker had been eliminated.

Concatenated SNPs were used to generate both maximum likelihood (ML) and Bayesian phylogenies without *a priori* assumptions, and with the ML analysis conducted in RAXML (v. 7.3.2; Stamatakis, 2006) using GTRCAT with 1,000 bootstraps. The Bayesian analysis was performed in MRBAYES (v. 3.2.3; Ronquist et al., 2012) using GTR (10,000,000 iterations), with sampling every 1,000 iterations and a 25% burn-in subsequently discarded.

However, methods employing concatenated SNPs can potentially overestimate support values for erroneous or poorly supported nodes (Liu et al., 2015; Edwards et al., 2016). This can be especially problematic in the presence of introgression, because the majority of loci may not support the resulting topology (Twyford & Ennos, 2012; Leaché et al., 2014a). However, multispecies coalescent methods perform well in situations where introgression is limited, and thus represent an important consideration when species with admixed ancestries are delimited (Edwards et al., 2016).

Applicability of these methods is limited with regards to SNP data, due to the common requirement of *a priori* inference of gene trees (see Leaché et al. 2017). Thus, multispecies coalescent inference was restricted to SVDquartets (Chifman & Kubatko, 2015) as implemented in PAUP* v. 4.0 (Swofford, 2003) that effectively bypasses the gene-tree inference step, thereby extending its applicability to SNP datasets. This approach uses a coalescent model to test support for quartets, and to calculate frequencies of SNPs for each species. The process does not require concatenation, but does necessitate the *a priori* partitioning of individuals into species or populations. Because of extensive run-times using exhaustive tip sampling, species were instead subdivided into populations based on high support under both concatenated SNP methods. All possible quartets were exhaustively sampled using 1000 bootstraps.

A multispecies coalescent phylogeny was generated from unlinked SNPs in SVDQUARTETS (Chifman & Kubatko, 2015) as implemented in PAUP* v. 4.0 (Swofford, 2003). This approach uses a coalescent model to test support for quartets, and to calculate frequencies of SNPs for each species. The process does not require concatenation, but does necessitate the *a priori* partitioning of individuals into species or populations. Species were subdivided into populations based on high support under both concatenated SNP methods. All possible quartets were exhaustively sampled using 1000 bootstraps.

### 2.5 Bayesian species delimitation

Species delimitation methods are a popular analytical approach, especially those coalescent-based (Fujita et al., 2012), and applicable to larger datasets. However, these can lead to over-splitting, particularly with respect to integrative taxonomic methods and Bayesian assignment tests (Miralles & Vences, 2013). The response of these methods to the effects of introgression are still tentative, and thus should be viewed with caution (Leaché et al., 2014b). Bayes Factor Delimitation (BFD; Leaché et al., 2014b) is another powerful tool for testing proposed taxonomic revisions, and to assess if models are congruent with the patterns of divergence obtained from multilocus genetic data. We applied it to test alternative models of species delimitation in: 1) Flannelmouth Sucker, 2) Bluehead Sucker, and 3) Zuni Bluehead Sucker. The latter is especially important, given the ongoing debate regarding its recent listing as an endangered subspecies (Federal Register, 2014).

BFD was performed using the SNP and AFLP Package for Phylogenetic analysis (SNAPP: Bryant et al., 2012). To accommodate assumptions and runtime limitations, we filtered the dataset to include only biallelic SNPs found across 95% of individuals, yielding data matrices of N=1,527 (FMS) and 1,742 (BHS). We estimated prior specifications for the population mutation rate (θ) as the mean pairwise sequence divergence within identified individuals of sister taxa (1.04×10^-3^ using *C. insignis* for FMS; 4.07×10^-4^ using *C. clarkii* for BHS). These were then used as the means for a gamma-distributed prior. We tested multiple prior-specifications for the lineage birth rate (λ) of the Yule model, using both fixed- and hyper-prior sampling of a gamma distribution. Fixed λ-values were calculated using PYULE (github.com/joaks/pyule), assuming tree height as ½ the maximum observed pairwise sequence divergence (i.e., 104.16 for FMS and 123.40 for BHS), and assuming the most conservative number of terminal nodes. Bayes factors for model comparison were calculated on normalized marginal likelihoods (Leaché et al., 2014b).

### 2.6 Hybrid detection

We calculated a hybrid index by mapping against interspecific heterozygosity. This served as a second means of assessing admixture, as well as to assess contemporary hybrid events. The est.h function (R-package INTROGRESS; Gompert & Buerkle, 2010) was used to estimate the hybrid index (Gompert & Buerkle, 2009) for samples at locations suggesting potential admixed ancestry. This included: 1) Rio Grande and Bluehead sucker (Zuni River, NM); 2) Sonora and Flannelmouth sucker [Little Colorado (AZ) and Virgin rivers (UT)]; and 3) Bluehead Sucker lineages (Little Colorado River). The calc.intersp.het and triangle.plot functions in INTROGRESS were also used to assess how contemporary were hybrid events. This was done by calculating interspecific heterozygosity, and by generating triangle plots for each admixture test, with recent hybrids identified according to high interspecific heterozygosity.

NEWHYBRIDS (Anderson & Thompson, 2002) was used to test the probability of hybrid assignment, to include first-filial (F1), second-filial (F2), first- and second-generation backcross (Bx), as well as those more ancestral (as gauged by Hardy-Weinberg expectations for random mating over several generations). Unlinked SNPs were used in both INTROGRESS and NEWHYBRIDS analyses, with additional filtering to remove loci that occurred: (a) Only in a single species, (b) In <80% of individuals, and (c) With a minimum allele frequency >10%.

## 3 Results

After filtering, a total of 20,928 loci and 98,230 SNPs were recovered in *Pantosteus,* with 60.8% (N=59,729) being parsimony-informative (PI) and 29.3% as missing data. For the subgenus *Catostomus*, 21,306 loci and 104,372 SNPs were recovered, with 66.4% (N=69,306) being PI, with 28.2% missing data. Unlinked SNPs (*Catostomus,* N=19,717; *Pantosteus,* N=20,038) were used to generate Bayesian clustering and multispecies coalescent phylogenies. Average coverage post-filtering was 17.8x, with all individuals at >8.9x coverage and with <80% missing data.

### 3.1 Phylogeny

Both concatenated SNP methods produced the same topology for each species (Figures 3A, 4A), with posterior probabilities of one and a bootstrap support of 100% for all species-level nodes, as well as for some populations within species. The multispecies coalescent phylogenies returned the same general topology as that produced by concatenated methods, but with variance in placement of the Rio Grande Sucker (*C. P. plebeius*)/ Desert Sucker (*C. P. clarkii*)/ Bluehead Sucker (*C. P. discobolus* and *C. P. virescens*) clade in the *Pantosteus* subgenus (Figure 3B). For the concatenated methods, Bluehead Sucker was placed outside the remaining species (Figure 3A), whereas for the multispecies coalescent method, Rio Grande Sucker was outside (Figure 3B). The latter reflects the results of previous research, to include morphological phylogenies, fossil evidence (Smith et al., 2013), as well as mitochondrial (Chen & Mayden, 2012; Unmack et al., 2014), and nuclear phylogenies (Bangs et al. 2018b). One impact of introgression was to obscure lineage-level topologies in *Catostomus* (Bangs et al. 2018b). Given this, we elected to emphasize full-lineage concatenated topologies, and note in so doing that topological discrepancies among the employed methods are both minimal, and reflective of processes that have been reviewed elsewhere.

**Figure 3:**
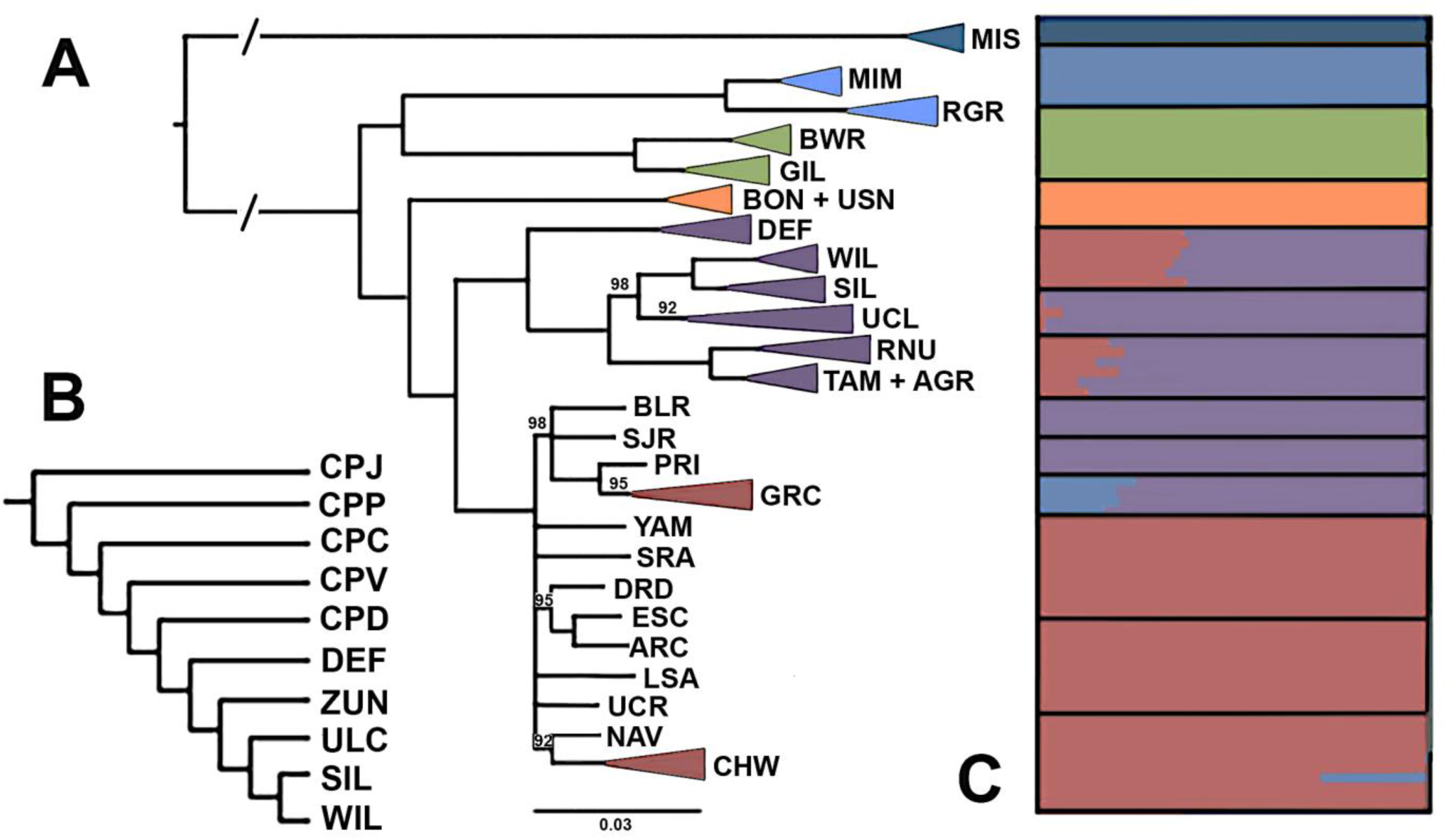
Phylogenetic and clustering results for subgenus *Pantosteus*. (A) Maximum likelihood phylogeny generated from 98,230 SNPs. Numbers represent bootstrap support, with nodes <80% collapsed. (B) Multispecies coalescent phylogeny generated from, 20,038 unlinked SNPs, with bootstrap support <100 indicated (CPJ= *Catostomus Pantosteus jordani*; CPP=*C.P. plebeius*; CPC=*C.P. clarki*; CVP=*C.P. virescens*; CPD=*C.P. discobolus*; DEF=Defiance Plateau; ZUN=Zuni River; ULC=Upper Little Colorado River; SIL=Silver Creek; WIL=Willow Creek); (C). Population clustering as provided by STRUCTURE, using 20,038 unlinked SNPs [arranged vertically as in (A)]. (MIS=Missouri River; MIM=Mimbres River; RGR=Rio Grande River; BWR=Bill Williams River; GIL=Gila River; BON=Bonneville Basin; USN=Upper Snake River; DEF=Defiance Plateau; WIL=Willow Creek; SIL=Silver Creek; UCL=Upper Colorado River; RNU=Rio Nutria; TAM=Tampico Springs; AGR=Agua Remora; BLR=Black Rocks; SJR=San Juan River; PRI=Price River; GRC=Grand Canyon; YAM=Yampa River; SRA=San Raphael; DRD=Dirty Devil; ESC=Escalante; ARC=Arch Canyon; LSA=Little Sandy; UCR=Upper Colorado River; NAV=Navajo; CHW=Chinle Wash).

For *Pantosteus*, isolated drainages were identified with high support in all phylogenetic analyses, to include: 1) Mimbres and Rio Grande rivers (Rio Grande Sucker), 2) Bill Williams and Gila rivers (Desert Sucker), and 3) Bonneville Basin, Upper Colorado and Little Colorado rivers (Bluehead Sucker) (Figure2, 3A, 3B). There was scant resolution among populations in the Upper Colorado River, but highly-supported nodes for MUs were consistent with those derived in previous microsatellite and mtDNA analyses (Hopken et al., 2013). Several highly supported groups were found within the Little Colorado River: 1) Defiance Plateau (AZ); 2) Willow Creek (AZ); 3) Silver Creek (AZ); 4) Upper Little Colorado River (AZ); and 5) Zuni River (NM) (Figures 2, 3A, 3B).

For *Catostomus*, highly supported splits were found not only between species but within Flannelmouth Sucker as well (i.e., Virgin, Upper Colorado, and Little Colorado rivers, Figure 4A). The Little Colorado River clade was sister to the Upper Colorado River, with the Virgin River outside of this grouping (Figure 4A, 4B). These results were consistent with previous phylogenomic results (Bangs et al. 2018b). ML analyses indicated three moderately-supported groups (80-90% bootstrap support) within the Little Colorado River: 1) Chevelon Canyon Lake (AZ); 2) Silver Creek (AZ); and 3) Wenima Wildlife Area (Upper Little Colorado River, AZ) (Figure 2, Figure 4A). All were supported at 1.0 Bayesian posterior probability in MRBAYES, but less so by SVDQUARTETS (<70% bootstrap support). Also, the split between Upper Colorado and Little Colorado rivers was only moderately supported (at 86%). It should be noted that Wenima was not included in the SVDQUARTETS phylogenetic analysis, due to hybridization with Sonora Sucker. However, its removal had no effect on topology or supports.

**Figure 4:**
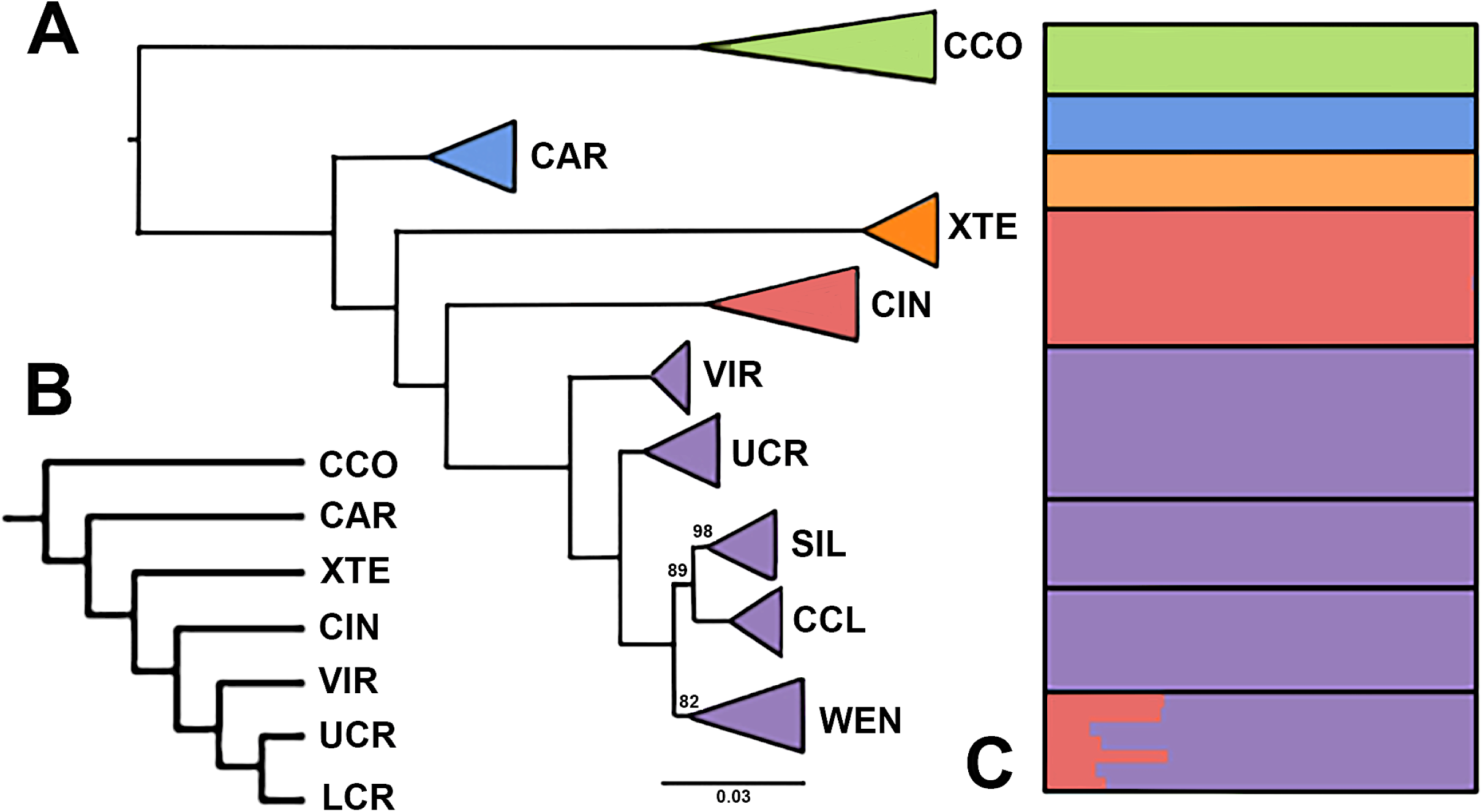
Phylogenetic and clustering results for subgenus *Catostomus*. (A) Maximum likelihood phylogeny generated from 69,306 SNPs. Numbers represent bootstrap support, with nodes <80% collapsed (CCO=*Catostomus commersonii*; CAR=*C. ardens*; XTE=*Xyrauchen texanus*; CIN=*C. insignis*; VIR=Virgin River; UCR=Upper Colorado River; LCR=Little Colorado River). (B) Multispecies coalescent phylogeny generated from19,717 unlinked SNPs, with bootstrap support <100 indicated (Abbreviations as in A, in addition to: SIL=Silver Creek; CCL=Chevelon Canyon Lake; WEN=Wenima Wildlife Area). (C). Population clustering as provided by STRUCTURE, using 19,717 unlinked SNPs [arranged vertically as in (A)].

### 3.2 Structure

The optimum number of supported clusters for *Pantosteus* was k=6, corresponding to: 1) Mountain Sucker (*C. P. jordani*); 2) Desert Sucker (*C. P. clarkii*); 3) Rio Grande Sucker (*C. P. plebeius*); and three clusters within Bluehead Sucker representing 4) Bonneville Basin (*C. P. virescens*); 5) Colorado River; and 6) Little Colorado River. The only Zuni River population assigned to Rio Grande Sucker was Rio Nutria (NM) (Figure 3C).

The only other mixing among *Pantosteus* clusters \was between Bluehead Sucker from the Colorado and Little Colorado rivers. This occurred in: 1) two (out of ten) samples from Chinle Wash (AZ), a tributary of the San Juan River; and 2) all Little Colorado River samples with the exception of Zuni River populations (the only group fully assigned to the Little Colorado River cluster). The proportion of assignments varied between regions in the Little Colorado River, but was largely consistent within each, with the Defiance Plateau (AZ) having the greatest assignment to the Colorado River cluster (32.7-38.6%), followed by populations from the Upper Little Colorado River (12.9-22.4%), and Willow and Silver creeks (AZ) (0.5-1.6%) (Figure 3C).

For *Catostomus*, the optimum number of supported clusters was k=5, corresponding to currently recognized species: 1) White Sucker (*C. commersonii*); 2) Utah Sucker (*C. ardens*); 3) Razorback Sucker (*Xyrauchen texanus*); 4) Sonora Sucker (*C. insignis*); and 5) Flannelmouth Sucker (*C. latipinnis*). No structure was apparent within Flannelmouth, even at higher k-values. Wenima was the only population to have mixed assignment, being allocated to both Flannelmouth and Sonora sucker, but with variation apparent in that four samples had lower assignments to Sonora Sucker (10.3-13.9%) when compared to the other three (26.9-28.3%). We interpret this as representing different hybrid classes (Figure 4C).

### 3.3 Hybridization

Individuals (N=4) from the Rio Nutria were tested for hybridization by employing Rio Grande Sucker (N=6) and Zuni Bluehead Sucker (N=8) as parentals. There were no missing data in the 302 unlinked SNPs [of which 59.2% (N=179) were fixed between species]. All four samples assigned with perfect support to the NEWHYBRIDS category “random mating over several generations.” An evaluation of Rio Nutria individuals by INTROGRESS yielded hybrid index values that were somewhat larger for Rio Grande Sucker (0.228-0.347) than the q-scores from STRUCTURE (0.170-0.252). However, the 95%-confidence intervals in INTROGRESS overlapped with the q-scores from STRUCTURE (Figure 5D), indicating agreement.

**Figure 5:**
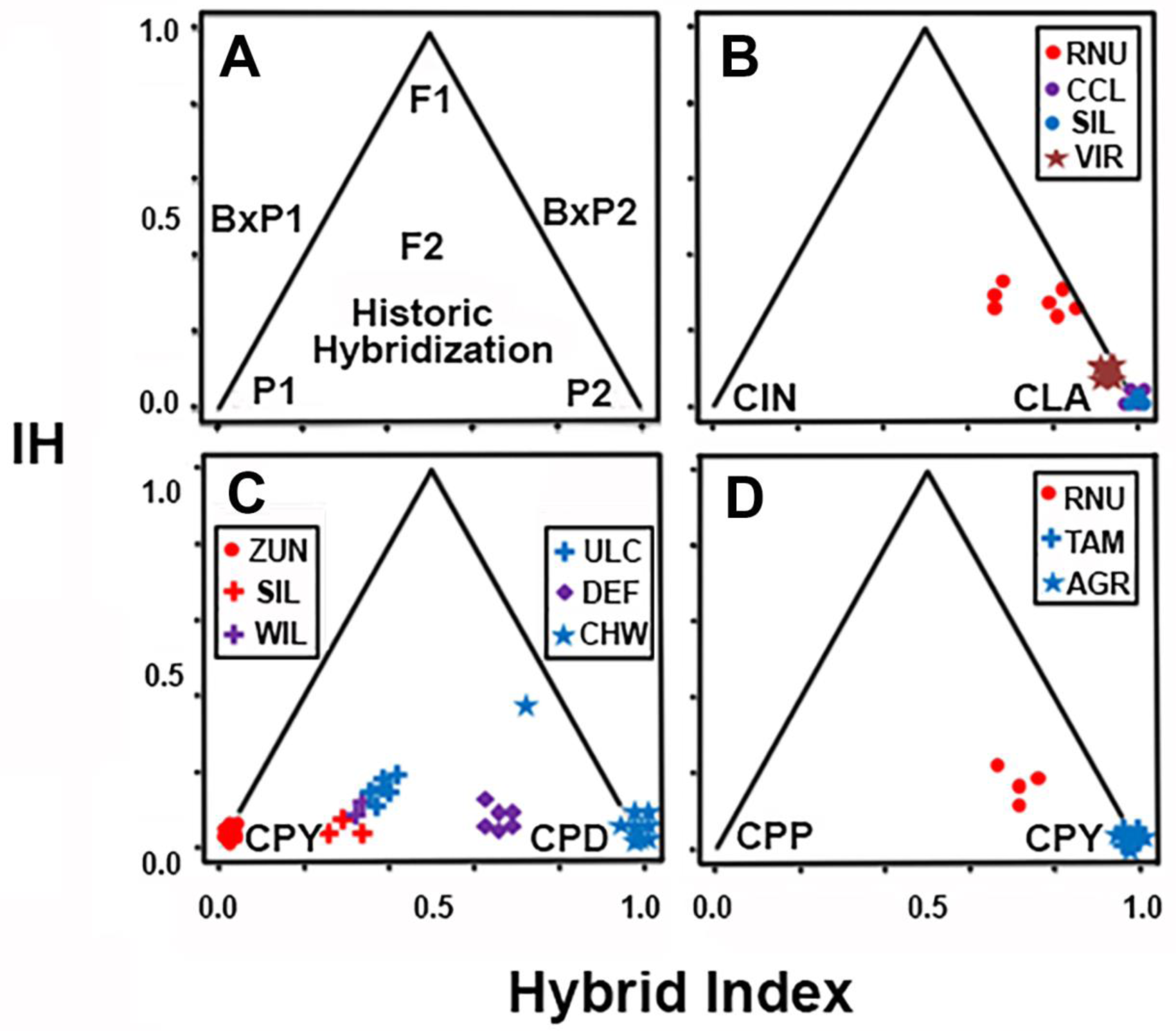
Triangle plots of interspecific heterozygosity versus hybrid index. (A) Hypothetical representation of pure, hybrid or backcrossed classes, with P1 and P2=Pure parental species; F1=First filial; F2=Second filial; and Bx=Backcross. (B) Sonora (CIN) x Flannelmouth sucker (CLA); (C) Zuni Bluehead (CPY) x Bluehead sucker (CPD); and (D) Rio Grande (CPP) x Zuni Bluehead sucker (CPY). Site abbreviations: AGR=Agua Remora; CCL=Chevelon Canyon Lake; CHW=Chinle Wash; DEF=Defiance Plateau; RNU=Rio Nutria; SIL=Silver Creek; TAM=Tampico Springs; ULC=Upper Little Colorado River; VIR=Virgin River; WEN=Wenima Wildlife Area; WIL=Willow Creek; ZUN=Zuni River.

Hybridization between Sonora and Flannelmouth sucker was also tested in the Little Colorado (N=14) and Virgin rivers (N=8), using Sonora Sucker (N=10) and the remaining Flannelmouth Sucker (N=13) as parentals. This analysis had 12.8% missing data, with 625 unlinked SNPs [of which 38.9% (N=243) were fixed between species]. Wenima was the only Little Colorado River population to reflect statistically significant hybridization with Sonora Sucker. An evaluation using INTROGRESS yielded slightly larger hybrid index values for Sonora Sucker (0.170-0.320) than the q-scores from STRUCTURE (0.103-0.283). Again, the 95% confidence interval generated by INTROGRESS overlapped with q-scores from STRUCTURE, indicating agreement. NEWHYBRIDS assigned four Wenima samples with greater than 95% probably as second generation (Bx) backcrosses into Flannelmouth Sucker. It also failed to assign the other three samples with high significance to any hybrid class, but instead assigned each to a mixture of different classes: F2, second generation backcrosses (Bx) into Flannelmouth Sucker, and “the random mating over several generations” category. All Flannelmouth Sucker samples from the Virgin River also had low, but significant, hybrid index values for Sonora Sucker (0.079-0.096). STRUCTURE did not detect this but it is consistent with the significant Patterson’s D-statistic of Bangs et al. (2018b) that point to historic introgression (Figure 5B).

Chinle Wash (N=10) and the Little Colorado River (N=17), with exclusion of Zuni River, were evaluated for mixing between the two clusters of Bluehead Sucker found in the Colorado River Basin. Here, parentals were: 1) Bluehead Sucker throughout the Upper Colorado River (N=21), and 2) those from Agua Remora and Tampico Springs of the Zuni River (N=8). The latter were used as they assigned completely to the Little Colorado River cluster in STRUCTURE (Figure 3C). A total of 546 unlinked SNPs served as input to INTROGRESS [17.9% fixed differences (N= 98) with 11.3% missing data]. Results essentially mirrored those of STRUCTURE, with the greatest hybrid index values for the Colorado River Bluehead cluster found in the Defiance Plateau (0.601-0.628), followed by Upper Little Colorado River (0.355-0.403), then Silver and Willow creeks (AZ) (0.266-0.333). Hybrid indices for all admixed individuals were significantly higher than q-scores, based on 95% confidence intervals in INTROGRESS. No overlap was found between the 95% CI of the hybrid index for any sample and the standard deviation between STRUCTURE runs. Two samples from Chinle Wash also showed significant admixture, with hybrid index values being 0.654 and 0.945 for the Colorado River cluster. The former had a high interspecific heterozygosity value, indicating potentially recent admixture (Figure 5C).

### 3.4 Bayes factor delimitation

To minimize the impact of introgression on species delimitation, all populations that showed significant introgression from outside species in STRUCTURE were removed from BFD runs. This included Rio Nutria (Bluehead Sucker) and Wenima (Flannelmouth Sucker), and included 1,527 and 1,742 unlinked SNPs respectively.

Splitting models were favored over lumping models in both Flannelmouth and Bluehead sucker, with the highest-ranked models being those with the most groups. Within Flannelmouth Sucker, the separation of the Virgin River was ranked higher than splitting either of the remaining two populations (i.e., Little Colorado and Colorado rivers). For Bluehead Sucker, splitting the Bonneville from the Colorado and Little Colorado rivers had greater ranking than splitting the Little Colorado River, but ranked lower than splitting all three. The currently-debated listings of the Zuni Bluehead Sucker, where the Zuni River or the Zuni-and-Defiance Plateau are split from the rest of the Little Colorado and Colorado rivers, were ranked lower than a model that split the Little Colorado River (to include Zuni River, Defiance Plateau, and Upper Little Colorado) from the Colorado River. However, the highest ranked model was one that split all three groups in the Little Colorado River (Table 2).

**Table 2:** Bayes Factor Delimitation for (A) Flannelmouth Sucker and (B) Bluehead Sucker. Group abbreviations under Model include: sos=Sonora Sucker, vir=Virgin River, lcr=Little Colorado River, col=Colorado River, des=Desert Sucker, bon=Bonneville Basin, zuni=Zuni River, def=Defiance Plateau, ulcr=Upper Little Colorado River (includes all populations above Grand Falls in the Little Colorado River, except for the Zuni River and Defiance Plateau). Marginal Likelihood (MarL) and Bayes Factor (BF) are shown for each model, with BF calculated by comparing to the least complex model for each species. Models are ranked by BF with the model most supported being above and least supported below. Models for Bluehead Sucker are split into two groups, with the last five involved with the splitting of the Zuni Bluehead Sucker. Results for BFD using alternative prior specifications did not vary, and so results only using a fixed λ are reported.

| A. Flannemouth Sucker |  |  |  |  |  |
| --- | --- | --- | --- | --- | --- |
| Model | Description of Model | # groups | MarL | BF | Rank |
| 1. sos, vir+lcr+col | Single species | 2 | -13009.53 | - | 5 |
| 2. sos, vir, lcr+col | Virgin River split | 3 | -12419.37 | -590.16 | 2 |
| 3. sos, lcr, vir+col | Little Colorado split | 3 | -12765.16 | -244.37 | 4 |
| 4. sos, col, vir+lcr | Colorado split | 3 | -12644.89 | -364.64 | 3 |
| 5. sos, vir, lcr, col | All split | 4 | -12334.89 | -674.64 | 1 |
| B. Bluehead Sucker |  |  |  |  |  |
| Model |  | # groups | MarL | BF | Rank |
| 1. des, bon+lcr+col | Single species | 2 | -24014.34 |  | 9 |
| 2. des, bon, lcr+col | *C. P. virecsens* split | 3 | -22678.56 | -2671.56 | 7 |
| 3. des, lcr, bon+col | Little Colorado split | 3 | -22758.92 | -2510.84 | 8 |
| 4. des, bon, lcr, col | Little Colorado + *C. P. virecsens* split | 4 | -21530.57 | -4967.54 | 4 |
| Zuni Bluehead Sucker Models |  |  |  |  |  |
| 5. des, bon, zuni, col+ulcr+def | Zuni River split | 4 | -21833.52 | -4361.64 | 5 |
| 6. des, bon, zuni+def, col+ulcr | Current listing on the Federal Register | 4 | -22178.81 | -3671.06 | 6 |
| 7. des, bon, col, zuni, ulcr+def | Zuni River split within Little Colorado | 5 | -20949.68 | -6129.32 | 2 |
| 8. des, bon, col, zuni+def, ulcr | Current listing with Little Colorado split | 5 | -21365.32 | -5298.04 | 3 |
| 9. des, bon, col, zuni, def, ulcr | All Little Colorado split | 6 | -20780.44 | -6467.8 | 1 |

## 4 DISCUSSION

Contemporary hybridization is problematic for freshwater conservation and management, particularly with regard to invasions (Bangs et al., 2018a; Hargrove et al., 2019) and translocations (Bruce & Wright, 2018). Yet these situations can most often be resolved through proper application of genomic approaches. However, it is much more difficult in a deep history context, in that phylogenetic relationships may be obscured as a result. Similarly, introgression is difficult to detect given genetic recombination (Wallis et al., 2016). Interestingly, freshwater fishes show particularly high levels of hybridization, due in large part to the occurrence of numerous sympatric species with small population numbers that are subsequently fragmented by environmental perturbations (Dowling & Secor, 1997). These issues have clearly impacted western North American freshwater fishes and, in particular, the genera evaluated herein (Dowling et al., 2016; Mandeville et al., 2017).

The six states that encompass the Colorado River Basin signed a ‘Range-wide Conservation Agreement Plan’ (2004) to adaptively manage our two study species basin-wide [as well as a third species, Roundtail Chub (*Gila robusta*)]. This, in turn, was a pre-emptive mechanism for these states to avoid potential listing under the Endangered Species Act (Carmen, 2007). All three species exhibit distinct life histories and habitat preferences that may have driven their potential divergences across the basin.

Since speciation is a gradual process with biodiversity elements scattered along its continuum (Sullivan et al., 2014), potential incongruence would be expected when different species delimitation methods are employed. Introgression would further complicate this process, yet its impacts on most species delimitation methods remain unknown, thus confounding any attempt to decipher results (Camargo et al., 2012). As such, the guideline of Carstens et al. (2013) are important considerations in this process, i.e., be conservative and employ multiple lines of evidence, given that a failure to delineate is expected. This includes the use of multiple algorithms for analyses of multi-locus data, and alternative lines of evidence that include (when possible) the life histories, morphologies, distributions, fossil histories, and behaviors of the biodiversity elements under study.

Here we explore different species delimitation approaches for two species, Flannelmouth and Bluehead sucker, to include the recent listing of the endangered Zuni Bluehead Sucker under the ESA. Our purpose was to evaluate similarities and differences in patterns of divergence in these two largely sympatric species with different life histories, and to diagnose (if appropriate) the potential for taxonomic revisions. In doing so, we also examined the impacts of introgression as a mechanism to disentangle their complex evolutionary histories that have evolved in lockstep with the geomorphology of the basin.

### 4.1 Life history and its effects on differentiation

Comparative phylogenomics of *Catostomus* and *Pantosteus* subgenera (per Smith et al., 2013) revealed parallel patterns throughout much of the Colorado River and neighboring basins (Bangs et al. 2018b). However, the scale of divergence varied greatly between these groups, as emphasized within the Upper Colorado River Basin (Figure 1).

Although three distinct clades were identified in Flannelmouth Sucker, they are relatively recent as underscored by the lack of distinct clustering (Figure 4) and having less than 1% fixed SNP sites as compared to >1.9% for all other comparisons (Table 3). These level of differentiation fits with recent events, including volcanic barriers that appeared during in the last 20kya, such as Grand Falls on the Little Colorado River (∼20kya).

**Table 3:** Total number of fixed SNPs between groups for (A) subgenus *Catostomus* and (B) subgenus *Pantosteus*. Below diagonal is the total number of fixed SNPs across all loci and above the diagonal is the proportion of fixed sites across all loci in percentage. In (A) the first three groups (Upper Colorado River Basin, Little Colorado River, and Virgin River) are Flannelmouth Sucker. In (B) the first three groups (Upper Colorado River Basin, Zuni River, and Bonneville Basin) are Bluehead Sucker.

|  | Upper<br>Colorado | Little<br>Colorado | Virgin River | Sonora<br>Sucker | Razorback<br>Sucker | Utah Sucker | White Sucker |
| --- | --- | --- | --- | --- | --- | --- | --- |
| Upper Colorado | - | 0.2% | 0.3% | 3.7% | 7.1% | 11.7% | 18.8% |
| Little Colorado | 216 | - | 0.9% | 4.2% | 7.1% | 10.9% | 16.7% |
| Virgin River | 314 | 986 | - | 4.1% | 8.1% | 13.3% | 20.2% |
| Sonora Sucker | 3879 | 4414 | 4263 | - | 7.6% | 12.9% | 19.3% |
| Razorback Sucker | 7381 | 7389 | 8493 | 7956 | - | 14.1% | 20.1% |
| Utah Sucker | 12249 | 11373 | 13837 | 13419 | 14719 | - | 15.4% |
| White Sucker | 19608 | 17445 | 21078 | 20102 | 21012 | 16090 | - |

B. Subgenus *Pantosteus*
|  | Upper<br>Colorado | Zuni River | Bonneville<br>Basin | Desert<br>Sucker | Rio Grande<br>Sucker | Mountain<br>Sucker |
| --- | --- | --- | --- | --- | --- | --- |
| Upper Colorado | - | 1.9% | 2.7% | 2.0% | 6.3% | 11.2% |
| Zuni River | 1862 | - | 5.7% | 6.2% | 9.4% | 13.1% |
| Bonneville Basin | 2611 | 5576 | - | 6.3% | 9.3% | 12.8% |
| Desert Sucker | 1985 | 6070 | 6231 | - | 7.3% | 12.1% |
| Rio Grande Sucker | 6176 | 9196 | 9181 | 7178 | - | 13.2% |
| Mountain Sucker | 11033 | 12908 | 12549 | 11917 | 12951 | - |

Lineages of Bluehead Sucker, on the other hand, reflect temporally deeper origins as underscored by the distinct clustering, branch lengths (Figure 3) and number of fix SNPs (1.9-3.3%; Table 3) that are equal to or greater than well-established species pairs represented in our analyses, as well as by previous mitochondrial dating (4.5-3.5mya; Unmack et al., 2014). However, the disentanglement of phylogenomic histories, and consequently the delineation of units for conservation and management, have been complicated by the secondary contact among lineages, as well as their hybridization with other species.

We suggest the contrasting timescales for these clades may stem from life history differences, particularly with regard to subgeneric habitat preferences. *Pantosteus* is commonly designated as ‘mountain sucker,’ due to its predilection for cooler habitats within higher elevation streams, whereas *Catostomus* is physically larger, omnivorous, and restricted to larger rivers that form lower-elevation components of basins (Sigler & Miller, 1962; Smith, 1966). Although Bluehead and Flannelmouth sucker largely co-occur, their habitat preferences are profound and must, in turn, influence diversification rates. For example, Douglas et al. (2003) suggested Flannelmouth Sucker in the Upper Colorado River Basin were driven into the Lower Basin by rapid Late Pleistocene warming and concomitant desiccation within the Upper Basin (e.g., the Hypsithermal; Pielou, 1974). It later recolonized the Upper Basin via the Grand Canyon. Although the same pattern was observed in mainstem Bluehead Sucker, populations likely persisted within the high elevation refugia that occurred in numerous tributaries of the Upper Colorado River Basin. This in turn would yield the shallow, but discernable genetic divergences among populations, and is consistent with the recognition of several as distinct management units (MUs) (Hopken et al., 2013).

### 4.2 Bonneville Basin

Although both species are sympatric in the Colorado River Basin, the Bluehead Sucker also occurs in the Bonneville and Upper Snake River basins (Figure 1). Therein, it may represent a unique species (originally described as *C. P. virescens* Cope & Yarrow 1875; Snyder, 1924) that was subsequently collapsed into *C. P. discobolus* (Smith, 1966). The split between *C. P. virescens* in the Bonneville Basin/ Snake River, and *C. P. discobolus* in the Colorado River Basin, is supported in all of our analyses. This includes population clustering, three different phylogenetic methods, and BFD analyses (Figure 3; Table 2). The convergence of all methods, along with recent morphological (Smith et al., 2013) and mitochondrial phylogenies (Hopken et al., 2013; Unmack et al., 2014), supports the reclassification of the Bonneville Bluehead Sucker. Furthermore, the chronology for the split between these two species (i.e., ∼4.8 mya per mtDNA time-calibrated phylogenies) exceeds that found in other catostomid species (Unmack et al., 2014), and emphasizes the deep divergence.

### 4.3 Little Colorado River Basin

Our phylogenetic analyses also separate the Little Colorado River Flannelmouth and Bluehead suckers from those in the Upper Colorado River Basin, to include the Grand Canyon (Figures 3, 4). The Little Colorado River lineages represent 1) Zuni Bluehead Sucker (*C. P. discobolus yarrowi*) now with a drastically reduced range that was influential in promoting its recent listing under the Endangered Species Act (Federal Register, 2014); and 2) Little Colorado River Sucker, currently recognized by Arizona Game and Fish Department as an undescribed species morphologically distinct from Flannelmouth Sucker (Miller, 1972; Minckley, 1980).

### 4.4 Zuni Bluehead Sucker

When *Pantosteus* was first described (Cope & Yarrow, 1875), the Zuni Bluehead Sucker was designated as a separate species. Subsequent allozymic and morphological data (Smith et al., 1983) not only recalibrated it to subspecies, but also suggested a hybrid origin that encompassed Bluehead and Rio Grande sucker. However, results from our studies now refute this hypothesis by demonstrating alleles from Rio Grande Sucker are found only within a single population (i.e., Rio Nutria) (Figure 3C, Figure 5D). This result is consistent with more contemporary analyses of allozymes (Crabtree & Buth, 1987) as well as single-gene sequencing data (Turner & Wilson, 2009; Hopken et al., 2013).

Zuni Bluehead Sucker seemingly originated in the mountains of northeast Arizona and northwest New Mexico, to include the Zuni River and Kin Lee Chee Creek of the Defiance Plateau (Smith et al., 1983). However, phylogenetic analyses render populations in Kin Lee Chee Creek and the Defiance Plateau as paraphyletic with the Zuni River and the remainder of the Little Colorado River (Figure 3A, 3B). In addition, the entire Little Colorado River Basin clade is a monophyletic group sister to the remainder of the Colorado River (Figure 3A, 3B). This suggests that Zuni Bluehead Sucker spread into the Little Colorado River following its integration with mountain streams (per Minckley, 1973; Smith et al., 1983). The current hypothesis (Smith et al., 1983) suggests that it was replaced by Bluehead Sucker in all Little Colorado River drainages, save Zuni River and Kin Lee Chee Creek.

However, population-clustering analyses (Figure 3C) yielded a clade unique to the Little Colorado River, within which only Zuni River populations were assigned. All other populations were assigned to a composite representing this cluster and the remainder of the Colorado River Basin, with proportions for the latter ranging from 0.5-38.6%. This admixture was also detected in hybrid index analyses, suggesting the remainder of the Little Colorado River Basin may represent an admixture of these two lineages (Figure 5C). Thus, Bluehead Sucker may have hybridized with Zuni Bluehead Sucker in the Little Colorado River rather than replacing it, with admixed populations now found in all but the Zuni River. The Defiance Plateau may be the source for this Bluehead Sucker invasion, based on a greater proportion of assignments with the Colorado River cluster. This may presumably be the result of stream capture with Chinle Wash (Figures 3C, 5C).

Further investigations employing a diversity of techniques (e.g., morphology, stable isotopes, and transcriptomes) may clarify how admixture has affected the breadth of lineages in the Little Colorado River. Our results support the Zuni Bluehead Sucker, and highlight the necessity of including the entire Little Colorado River clade when its status is assessed. This is particularly highlighted in the model testing of BFD, where the current listing (to include both the Zuni River and Kin Lee Chee Creek of the Defiance Plateau) was ranked lower than either a splitting of the entire Little Colorado River, or just the Zuni River (Table 2). This necessitates a reassessment of the Zuni Bluehead distribution, so as to either separate from it the Kin Lee Chee Creek population or include it within the Little Colorado River Basin.

### 4.5 Little Colorado River Sucker

In contrast to the Zuni Bluehead Sucker, the Little Colorado River Sucker did not cluster separately, despite its representation as a monophyletic group in all phylogenetic analyses (Figure 4). This may reflect its recent origin, concomitant with formation of Grand Falls ∼20kya. This vicariant break effectively separated the Upper Little Colorado River from the rest of the Colorado River, and prevented contemporary upstream gene flow (Duffield et al., 2006). Although similar contemporary phylogeographic patterns are found in Zuni Bluehead Sucker and Little Colorado River Sucker, different evolutionary histories are apparent, as driven by habitat preference. This process ultimately resulted in levels of divergence that differ, but within similar contemporary ranges. This underscores the chaotic fluvial history of the Desert Southwest, as well as the need for comparative studies that can disentangle the organismal histories that coexist there.

Hybridization was also detected between Sonora and Flannelmouth sucker in Wenima Wildlife Area of the Little Colorado River (Figure 4C). These admixed individuals are presumably due to a recent hybrid event, as gauged by the variation in q-scores found in Sonora Sucker (Figure 4C), as well as hybrid index values (Figure 5B), high interspecies heterozygosity (Figure 5B), and the presence of four second-generation hybrids. Regardless, further sampling is needed to confirm this assumption.

### 4.6 Virgin River

Despite forming a monophyletic group, the Little Colorado River Sucker fell within a paraphyletic Flannelmouth Sucker. This was due largely to the placement of the Virgin River (Figure 4), also suggested as potentially unique due to an elevated morphological variability stemming from potential hybridization with Sonora Sucker (*C. insignis*) and Razorback Sucker (*Xyrauchen texanus*) (Minckley, 1980). Indeed, historic introgression with Sonora Sucker was detected in all Virgin River samples, as reflected in the elevated hybrid index and low interspecies heterozygosity (Figure 5B). Although the Sonora Sucker proportion is reduced, it is nevertheless significant based on previous D-statistic tests (Bangs et al., 2018b) and hybrid index values for all samples (Figure 5B).

Although the phylogenetic splitting of the three Flannelmouth Sucker groups (i.e., Upper Colorado, Little Colorado, and Virgin River) was also supported in BFD (Table 2). they grouped as a single cluster in STRUCTURE (Figure 4C) and the splits could not be replicated in cluster analyses, even at higher k-values. This, in turn, suggests a recent origin for these groups, further supported by their short branch lengths (Figure 4A). There is also a lack of fixed differences between these lineages in a previous mitochondrial analysis (Douglas et al., 2003). These considerations fit well with the previous assumption that the Virgin River population may have separated recently, i.e., Late Pleistocene, most likely due to climatic oscillations that alternately connected and separated Grand Canyon and Virgin River as recently as 7.5kya (Douglas et al., 2003). The support in BFD for the splitting of these groups may be due to an increased sensitively in defining recent splits, or may instead be biased by differential introgression with Sonora Sucker, particularly given the unknown capacity of this method to discern introgression (Leaché et al., 2014b).

## 5 CONCLUSIONS

Flannelmouth and Bluehead sucker are recognized as ‘species of concern’ in the Colorado River Basin (Carmen, 2007). Proposed taxonomic revisions will not only impact the management of these species, but also the basin as a biogeographic unit. Both species reflect similar phylogenomic patterns, yet their levels of divergence underscore evolutionary histories that differ significantly, and which impact their species delimitations. Three lineages of Bluehead Sucker were detected in all phylogenetic and population genetic methods, with *C. P. virescens* in the Bonneville and Upper Snake River elevated as a species separate from *C. P. discobolus* in the Colorado River (per Smith et al., 2013; Unmack et al., 2014). Results also support the Zuni Bluehead Sucker as a unique form. However, the current designation of Kin Lee Chee Creek as congruent with the Zuni River is erroneous, as they are instead paraphyletic. This situation can be resolved by including the of Little Colorado River Bluehead Sucker, or by removing Kin Lee Chee Creek from the listing of the Zuni Bluehead Sucker under the ESA. The situation is further complicated by hybridization with Rio Grande Sucker, and Bluehead Sucker from the Colorado River.

The Little Colorado River Sucker falls within a paraphyletic Flannelmouth Sucker, and can only be resolved by designating the Virgin River population as a unique lineage. However, these three clades are of recent origin, based on population genetic analyses (herein) and the lack of resolution found in mitochondrial analyses (Douglas et al., 2003). Thus, all three Flannelmouth Sucker lineages are more accurately represented as evolutionary significant units (ESUs), as reflected by their reduced phenotypic and genetic differentiation. They thus lack concordance under the genealogical component of the phylogenetic species concept.

## ACKNOWLEDGEMENTS

Numerous agencies contributed field expertise, specimens, technical assistance, collecting permits, funding or comments for the completion of this study. We are in debt to students, postdoctorals, and faculty who contributed to the development of our research: S. Mussmann, A. Albores, P. Brunner, M. Davis, T. Dowling, R. Cooper, J. Cotter, E. Fetherman, M. Hopken, M. Kwiatkowski, A. Reynolds, J. Reynolds, C. Secor, and P. Unmack. Sampling procedures were approved under IACUC permit 98-456R (Arizona State University) and 01-036A-01 (Colorado State University). Samples were also provided by the Museum of Southwestern Biology, University of New Mexico (T. Turner, L. Snyder). Accession numbers for samples from the Museum of Southwestern Biology are: MSB 44046, 44047, 44052, 44059, 44067, 49715, 49962, 53500, 63249, and 88362. Analyses were supported by the Arkansas High Performance Computing Center (AHPCC), funded through multiple National Science Foundation grants and the Arkansas Economic Development Commission. This research was also supported by generous endowments to the University of Arkansas: The Bruker Professorship in Life Sciences (M.R. Douglas), and 21st Century Chair in Global Change Biology (M.E. Douglas).

## AUTHOR CONTRIBUTIONS

MRB, MRD and MED designed the study; MRB prepared DNA and generate ddRAD libraries; MRB and TKC completed data analyses; all authors contributed in drafting the manuscript and all approved its final version.

## DATA ACCESSIBILITY

Raw fastq files for each individual as well as all alignments used in this study are available on the Dryad Digital Repository (address added after acceptance of article).

